# Glucocorticoid-Induced Metabolic Disturbances are Exacerbated in Obesity

**DOI:** 10.1101/184507

**Authors:** Innocence Harvey, Erin J. Stephenson, JeAnna R. Redd, Quynh T. Tran, Irit Hochberg, Nathan Qi, Dave Bridges

## Abstract

Objective: To determine the effects of glucocorticoid-induced metabolic dysfunction in the presence of diet-induced obesity. Methods: C57BL/6J adult male lean and diet-induced obese mice were given dexamethasone for different durations and levels of hepatic steatosis, insulin resistance and lipolysis were determined. Results: Obese mice given dexamethasone had significant, synergistic effects on insulin resistance and markers of lipolysis, as well as hepatic steatosis. This was associated with synergistic transactivation of the lipolytic enzyme ATGL. Conclusions: The combination of chronically elevated glucocorticoids and obesity leads to exacerbations in metabolic dysfunction. Our findings suggest lipolysis may be a key player in glucocorticoid-induced insulin resistance and fatty liver in individuals with obesity.

## Introduction

Cushing’s syndrome manifests in response to chronically elevated levels of glucocorticoids and is often associated with changes in adipose mass and distribution, non-alcoholic fatty liver disease (NAFLD) and impaired glucose tolerance (Paredes & Ribeiro 2014). While Cushing’s disease is rare, it is estimated that at any given time 1-3% of the US, UK and Danish populations are prescribed exogenous corticosteroids, which may increase their risk for developing the metabolic complications (Hsiao *et al.* 2010; Fardet *et al.* 2011; Overman *et al.* 2013; Laugesen *et al.* 2017).

Similarly, obesity is accompanied by a multitude of metabolic disturbances, such as insulin resistance and NAFLD and is a worldwide epidemic (The GBD 2015 Obesity Collaborators 2017). Comparing the high rates of medically prescribed corticosteroids with the prevalence of overweight and obesity in developed countries, the combination of obesity and glucocorticoid excess may be present in many individuals. Given the similar co-morbidities associated with obesity and chronically elevated glucocorticoids, we hypothesized that the combinations of these two conditions would lead to worse metabolic outcomes than either of them alone. This is supported by studies in rats showing that corticosterone and high-fat diets combine to cause worsened insulin resistance and non-alcoholic fatty liver disease (Shpilberg *et al.* 2012; Beaudry *et al.* 2013).

There is an array of physiological changes that occur as a result of elevated glucocorticoids including decreased lean mass (Dardevet *et al.* 1995; Schakman *et al.* 2013; Hochberg *et al.* 2015), increased fat mass (Abad *et al.* 2001; Geer *et al.* 2011; Hochberg *et al.* 2015), NAFLD (Shpilberg *et al.* 2012) and increased lipolysis (Djurhuus *et al.* 2002, 2004; Kršek *et al.* 2006), all of which have been associated with decreased insulin sensitivity (Rebrin *et al.* 1996; Zhang *et al.* 2015; Dirks *et al.* 2016). Recent tissue-specific knockouts of glucocorticoid signaling mediators have implicated adipose tissue as a central node linking glucocorticoid action and lipolysis to systemic insulin resistance and NAFLD (Morgan *et al.* 2014; Wang *et al.* 2014; Mueller *et al.* 2017; Shen *et al.* 2017). Here we present the finding that chronically elevated glucocorticoids in the presence of diet-induced obesity have synergistic effects on lipolysis, insulin resistance and fatty liver disease. Obese glucocorticoid-treated mice have reduced fat mass compared to all other groups, yet have hyperglycemia and severe insulin resistance. Therefore, we speculate that lipolysis drives insulin resistance in obese animals.

## Methods

### Animal Procedures

C57BL/6J adult male mice were purchased from the Jackson Laboratory at nine weeks of age (stock #000664). All animals were on a light dark cycle of 12/12 h and housed at 22°C. Following a week of acclimation, mice were placed on diets or treated with dexamethasone as described in the figure legends. Mice were treated with vehicle (water) or approximately 1mg/kg/d of water-soluble dexamethasone (Sigma-Aldrich; catalog #2915) dissolved in their drinking water for 12 weeks, as described previously (Hochberg *et al.* 2015). Additional cohorts of mice used in these experiments either remained on a standard diet (normal chow diet; NCD; 5L0D LabDiet; 13% fat; 57% carbohydrate; 30% protein) or were provided a high fat diet (45% fat from lard; 35% carbohydrate mix of starch, maltodextrin and sucrose; 20% protein from casein; cat# D12451) for either 8 or 12 weeks followed by dexamethasone treatment. Mice were group housed with four mice per cage and food consumption was measured weekly by weight reductions per cage and calculated to reflect estimated intake of each mouse per day in a given cage. Mice remained on their respective diets for the duration of the study. All mice were provided with access to food and water *ad libitum* throughout the study, unless otherwise noted. For the longer, six-week dexamethasone treatments, 16 HFD-fed, dexamethasone-treated mice appeared ill and were euthanized and thus removed from all analyses once symptoms were noticed. Symptoms included lethargy, weight loss and evidence of pancreatitis in some of the mice. Animal body weight and composition was determined weekly using a digital scale and EchoMRI 2100, respectively. We performed a CLAMS experiment (data not shown) with the 12-week diet study prior to dexamethasone treatment where mice were singly housed for approximately one week, which led to fluctuations in body weight over the first week. Body weight quickly stabilized following removal from the CLAMS in both groups. At the end of treatment, mice were fasted for 16h, dexamethasone water was not removed during this time, and euthanized by cervical dislocation after isoflurane anesthesia at ZT3. Immediately following euthanasia, mice were dissected and the right inguinal white adipose tissue (iWAT) and epididymal white adipose tissue (eWAT) depots were carefully removed and weighed. Adipose tissues, along with a section of the left lateral lobe of the liver were snap frozen in liquid nitrogen for later analysis. Small pieces of tissues were fixed in 10% phosphate-buffered formalin for histology. Animal procedures were approved by the University of Tennessee Health Science Center and University of Michigan Institutional Animal Care and Use Committees.

### Insulin Tolerance Tests and Hyperinsulinemic Euglycemic Clamp Experiments

Insulin responsiveness was assessed via an insulin tolerance test (ITT). Following a six hour fast, mice were given an intraperitoneal (IP) injection of insulin (Humulin R, Lilly,) as described in figure legends. Blood was collected from the tail and glucose was determined using a One Touch Ultra Glucometer (Lifescan). For the hyperinsulinemic euglycemic clamp experiments, C57BL/6J adult (70d) male mice were fed HFD for eight weeks and treated with dexamethasone in their drinking water for three weeks or regular drinking water. Animals were anesthetized with an IP injection of sodium pentobarbital (5060 mg/kg). Indwelling catheters were inserted into the right jugular vein and the right carotid artery, respectively. The free ends of catheters were tunneled subcutaneously and exteriorized at the back of the neck via a stainless-steel tubing connector (coated with medical silicone) that was fixed subcutaneously upon closure of the incision. Animals with healthy appearance, normal activity, and weight regain to or above 90% of their pre-surgery levels were used for the study. Experiments were carried out in conscious and unrestrained animals using techniques described previously (Halseth *et al.* 1999; Ayala *et al.* 2006; McGuinness *et al.* 2009). Briefly, the primed (1.0 *μ*Ci)-continuous infusion (0.05 *μ*Ci/min and increased to 0.1 *μ*Ci/min at t = 0) of [3-^3^H] glucose (50 *μ*Ci/ml in saline) was started at t = -120min. After a five-hour fast, the insulin clamp was initiated at t = 0, with a prime-continuous infusion (40 mU/kg bolus, followed by 8.0 mU/kg/min) of human insulin (Novo Nordisk). Euglycemia (120~30 mg/dL) was maintained during the clamp by measuring blood glucose every 10 min and infusing 50% glucose at variable rates, accordingly. Blood samples were collected from the right carotid artery at t = 80, 90, 100, and 120 min for determination of glucose specific activity. Blood insulin concentrations were determined from samples taken at t = -10 and 120 min. A bolus injection of [1-^14^C]-2-deoxyglucose ([^14^C]2DG; PerkinElmer) (10 *μ*Ci) was given at t = 120 min. Blood samples were taken at 2, 5, 10, 15, and 25 min after the injection for determination of plasma [^14^C]2DG radioactivity. At the end of the experiment, animals were anesthetized with an intravenous injection of sodium pentobarbital and tissues were collected and immediately frozen in liquid nitrogen for later analysis of tissue [1-^14^C]-2-deoxyglucose phosphate ([^14^C]2DGP) radioactivity. Blood glucose was measured using an Accu-Chek glucometer (Roche, Germany). Plasma insulin was measured using the Linco rat/mouse insulin ELISA kits. For determination of plasma radioactivity of [3-^3^H]glucose and [1-^14^C]2DG, plasma samples were deproteinized with ZnSO_4_ and Ba(OH)_2_ and counted using a Liquid Scintillation Counter (Beckman Coulter LS6500 Multi-purpose Scintillation Counter). Glucose turnover rate, hepatic glucose production and tissue glucose uptake were calculated as described elsewhere (Kraegen *et al.* 1985; Halseth *et al.* 1999; Ayala *et al.* 2006).

### Serum Glycerol and Fatty Acid Determination

Following 12 weeks of dex-amethasone treatment, 22-week-old *ad libitum* chow fed C57BL/6J male mice were anesthetized with isoflurane and blood was collected into heparin-coated capillary tubes via retro orbital bleed both prior to and 15 minutes following intraperitoneal injection of 10mg/kg isoproterenol (Sigma-Aldrich; catalog #I6504-1G) in Dulbecco’s phosphate-buffered saline (Thermo Fisher; catalog #BW17512F1). Serum from these mice, as well as from a cohort of 28-week old mice on either HFD or chow, six-weeks post-dexamethasone treatment was collected following an overnight fast. Glycerol was assessed via Serum Triglyceride Determination Kit (Sigma-Aldrich; catalog #TR0100-1KT) and fatty acids were quantified using the HR Series NEFA-HR(2) kit (Wako Diagnostics; catalog #276-76491), in accordance with manufacturer’s guidelines.

### Cell culture

3T3-L1 fibroblasts (pre-adipocytes; ATCC; authenticated via STRS analysis) were cultured in 10% newborn calf serum, Dulbecco’s Modification of Eagle’s Medium (DMEM; 4.5 g/L D-glucose; Fisher Scientific; catalog #11965118) with penicillin, streptomycin and glutamine (PSG) until confluence. Cells were switched to a differentiation cocktail at two days post confluence (250nM dexamethasone, 500*μ*M 3-isobutyl-1-methylxanthine and 1*μ*g/mL insulin in 10% fetal bovine serum, in 4.5g/L glucose DMEM with PSG) for four days (Chiang S-H, Chang L 2002). Media was replaced with differentiation medium containing only insulin for an additional three days. For the following three days, cells remained in media with no additional treatment. Cells used for these experiments were not cultured beyond 22 passages. To assess markers of lipolysis, cells remained in media and were treated with ethanol (vehicle) or 250nM dexamethasone for five days before lysing.

### Assessment of Triglyceride Content in Cells and Tissue

3T3-L1 cells were grown and treated as described above. At the end of the treatment period, cells were lysed in homogenization buffer (50 mM Tris pH 8, 5 mM EDTA, 30 mM Mannitol, protease inhibitor) and subjected to three freeze thaw cycles with liquid nitrogen, thawed at room temperature. Frozen liver tissue was homogenized using a TissueLyser II (Qiagen). Lipids were extracted using KOH and a chloroform to methanol (2:1) extraction. Triglyceride content was assessed using the Serum Triglyceride Determination Kit and absorbance was detected as described in (Lu *et al.* 2014).

### Histology

Tissues were fixed in 10% phosphate-buffered formalin for 24 hours and then stored in 70% ethanol until further processing. Tissues were dehydrated, embedded in paraffin and sent to the University of Michigan Comprehensive Cancer Center Tissue Core where they were processed and stained with hematoxylin and eosin (H&E) to assess cell morphology. Slides were imaged using the 10x objective of an Olympus iX18 inverted microscope and cellSense software.

### mRNA Extraction and Analysis

Cells and tissues were lysed in TRIzol using the TissueLyser II, as decribed above, and RNA was extracted using a Pure-Link RNA kit (Life Technologies; catalog #12183025). cDNA was synthesized from 0.5–1*μ*g of RNA using the High Capacity Reverse Transcription Kit (Life Technologies; catalog #4368813). Primers, cDNA and Power SYBR Green PCR Master Mix (Life Technologies; catalog #4368708) were combined in accordance with the manufacturer’s guidelines and quantitative real-time PCR (qPCR) was performed as previously described (Lu *et al.* 2014) using the QuantStudio 5 (Thermo Fisher Scientific). mRNA expression level was normalized to *Actb* and analyzed using the ΔΔ Ct method after evaluation of several reference genes. qPCR primer sequences are listed in Table 1.

**Table 1:**
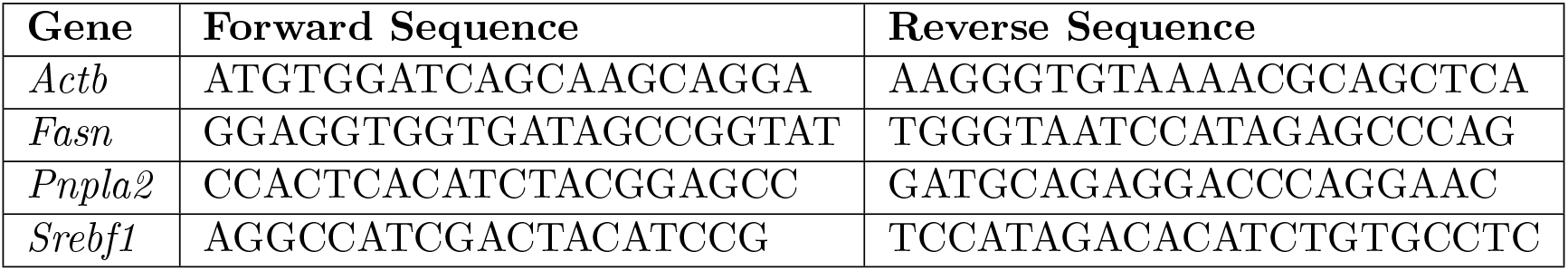
Primers used for RT-qPCR analyses.

### Protein Extraction and Analysis

Cells and tissues were lysed in RIPA buffer (50 mM Tris, pH 7.4, 0.25% sodium deoxycholate, 1% NP40, 150 mM sodium chloride, 1 mM EDTA, 100 uM sodium orthovanadate, 5 mM sodium fluoride, 10 mM sodium pyrophosphate and 1x protease inhibitor), centrifuged at 14,000rpm for 10 minutes at 4^°^C. Lysates were heated with loading buffer at 85–95^¤^C and proteins were separated by SDS-PAGE (Life Technologies) and transferred onto nitrocellulose membranes overnight at room temperature. Membranes were blotted at room temperature using anti-adipose triglyceride lipase antibodies (ATGL; 54 kDa; Cell Signaling Technologies; catalog #30A4). Antibody complexes were detected by anti-mouse and anti-rabbit fluorescent conjugated antibodies (Invitrogen) and visualized using an Odyssey CLx image scanner. Blots were quantified using Image Studio software version 5.2 (LiCOR) and normalized to Revert Total Protein Stain (LiCOR; catalog #926-11011).

### Statistics

All data are presented as mean +/- standard error of the mean. For animal studies, two-way ANOVA analyses were performed to test for significance of diet and dexamethasone treatment, as well as their interaction. Pairwise comparisons, normality and equal variance were tested using Shapiro-Wilk and Levene’s tests, respectively. Pending those results, a Mann-Whitney, Welch’s or Student’s *t*-test were used. P-values below p=0.05 were considered significant. All statistical tests were performed using the R software package version 3.30. All raw data and analysis scripts are available at http://bridgeslab.github.io/CushingAcromegalyStudy/.

## Results

### Dexamethasone-Induced Insulin Resistance is Worsened in the Presence of Obesity

To investigate if obesity status influences insulin sensitivity in the presence of elevated glucocorticoids we performed an insulin tolerance test (ITT) on lean (NCD) and diet-induced obese (HFD) mice that were untreated (water) or treated with glucocorticoids (dexamethasone). HFD-fed, dexamethasone-treated mice were significantly more resistant to insulin-stimulated glucose disposal when compared to all other groups (Figure 1A). Additionally, HFD dexamethasone-treated mice exhibited dramatic fasting hyperglycemia, with a significant interaction between diet and drug (p=0.00009; Figure 1B). While HFD animals had a 24% increase in fasting glucose when compared to NCD animals, in the presence of dexamethasone, HFD-fed animals had a 122% increase in fasting glucose relative to NCD controls not treated with dexamethasone. In the lean, NCD-fed animals, dexamethasone caused an 18% decrease in fasting glucose.

**Figure 1:**
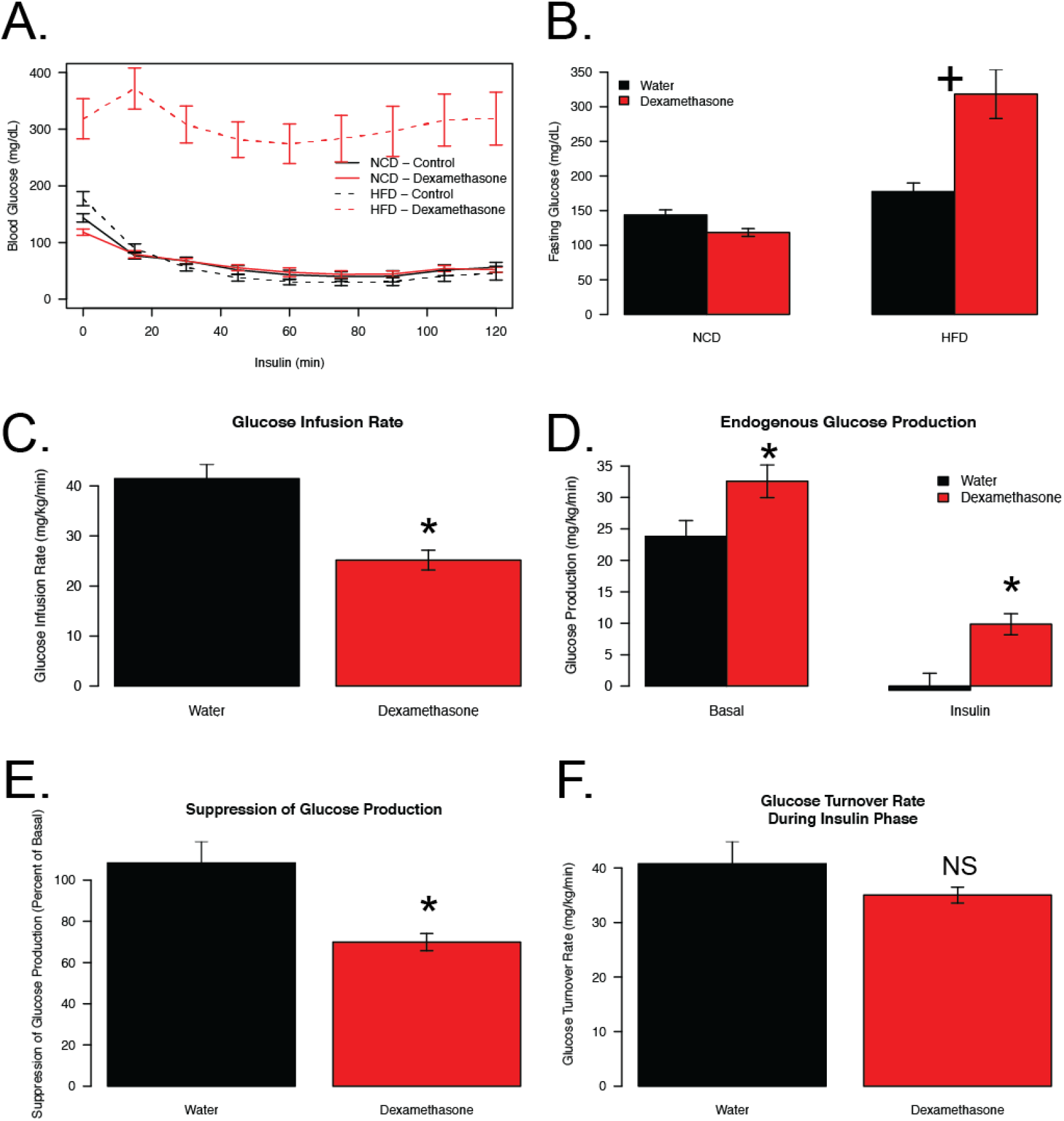
Reductions in glucose handling are exacerbated when elevated glucocorticoids and obesity are combined. Mouse blood glucose levels during insulin tolerance test (C) and prior to insulin injection (basal; D). Insulin was given via i.p. injection at a concentration of 2.5 U/kg following five weeks of dexamethasone (NCD n=12; HFD n=12) or vehicle (NCD n=12; HFD n=12) treatment and 17 weeks of diet. Mouse glucose infusion rate (GIR; E) and endogenous glucose production (EGP; F) during euglycemic clamp following 3 weeks of dexamethasone (n=14) or vehicle (n=11) treatment and 11 weeks of HFD. For clamp experiments, insulin was infused at 8 mU/kg/min following a prime continuous infusion of 40mU/kg bolus. All mice were fasted for 5-6 hours prior to experiments. Crosses indicate a significant interaction between diet and treatment. Asterisks indicate a statistically significant treatment effect for the pairwise comparison.

To evaluate glucose homeostasis in more detail we performed hyperinsulinemic-euglycemic clamps in obese mice (11 weeks of HFD) treated with dexamethasone for the final three weeks. This shorter HFD/dexamethasone exposure still caused dramatic insulin resistance, hyperglycemia and reductions in lean mass (Supplementary Figures 1A-D). Animals were clamped while conscious and glucose levels during the clamp as well as insulin turnover rate were similar between groups (Supplementary Figure 1E,F). During the hyperinsulinemic phase, the glucose infusion rate was 39% lower in obese dexamethasone-treated mice when compared to obese controls indicating insulin resistance at euglycemia (Figure 1C). Basal endogenous glucose production (EGP) was 37% higher in the dexamethasone-treated group (p=0.026). Moreover, in the control group, EGP was reduced to near zero by a high dose of insulin but only reduced 70% in the dexamethasone group (p=0.0091) resulting in glucose production being higher during the insulin phase in dexamethasone-treated mice (p=0.014) when compared to controls (Figure 1D-E). Glucose turnover was slightly decreased in the presence of insulin (p=0.141; Figure 1F). Despite these modest changes in glucose turnover, there were significant reductions in the obese, dexamethasone-treated animals in 2-deoxyglucose uptake in heart (34% reduced, p=0.0003) and gastrocnemius tissues (68% reduced; p=0.00002; Supplementary Figures 1G-H). These data suggest that increased glucose production and its impaired suppression by insulin are the likely causes of poor glycemic control in obese, dexamethasone-treated animals.

### HFD-Induced Liver Steatosis in Dexamethasone-Treated mice

Obesity and chronic elevations in glucocorticoids are associated with NAFLD (Wanless & Lentz 1990; Rockall *et al.* 2003). H&E staining of hepatic tissue clearly depicts exacerbated lipid levels in the obese, dexamethasone-treated group when compared to obese controls and lean groups (Figure 2A). In support of this, we observe drastically elevated liver triglycerides when compared to all other groups with a significant interaction between drug and diet (p=0.000068; Figure 2B).

**Figure 2:**
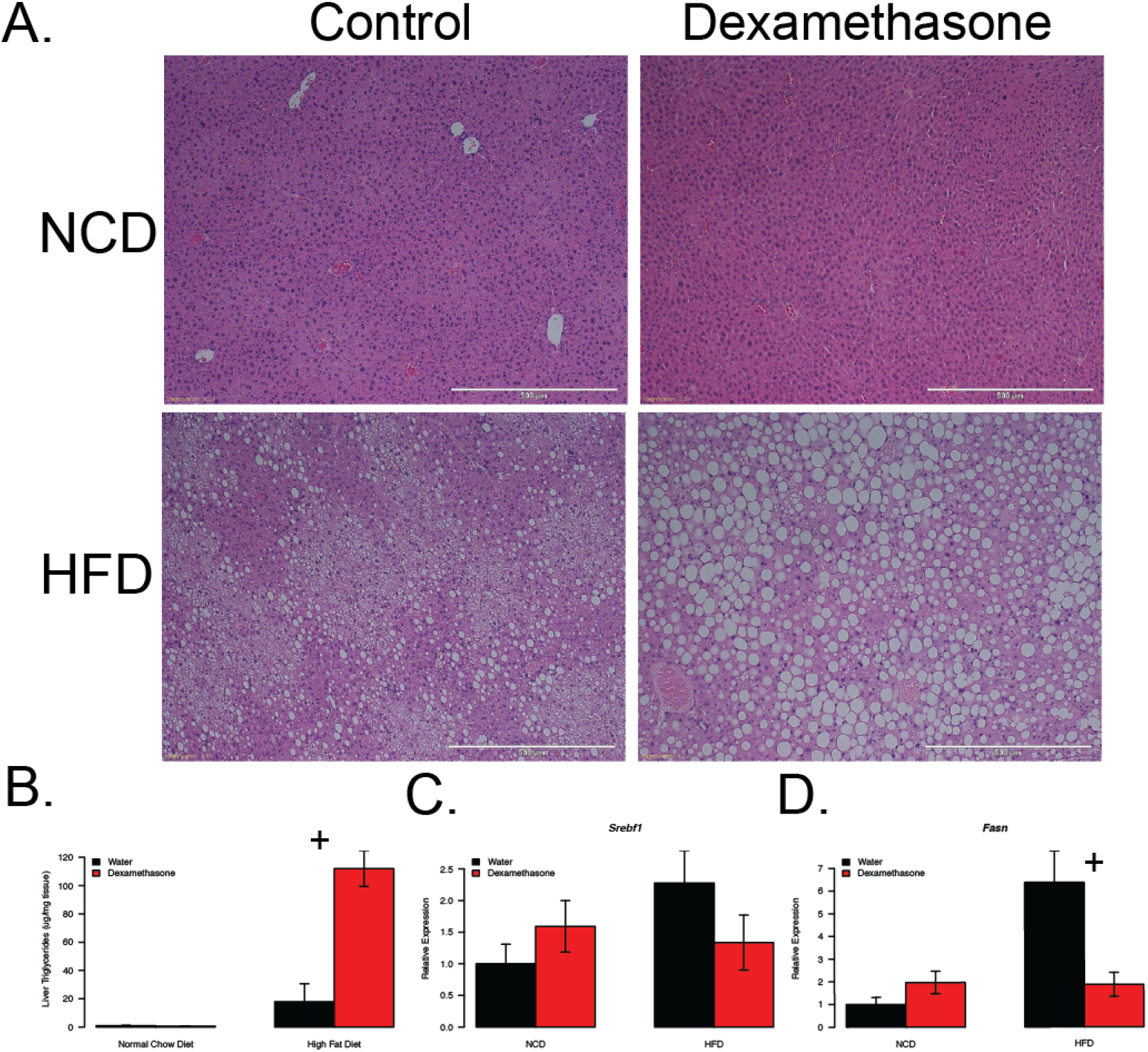
Increased glucocorticoids lead to greater severity of hepatic steatosis in obese mice. Mouse hepatic triglyceride levels (B) and Hematoxylin and Eosin stained liver sections (C) and qPCR of hepatic *de novo* lipogenic transcripts (D, E). Mice were euthanized at 28 weeks of age following six weeks of dexamethasone (NCD n=7; HFD n=5) or vehicle (NCD n=6; HFD n=9) treatment and 18 weeks of diet. Liver stains are representative samples from each group. Crosses indicate a significant interaction between diet and treatment.

We used qPCR to measure the expression of genes involved in hepatic *de novo* lipogenesis, *Srebf1* and *Fasn,* in liver lysates (Figure 2C-D). We observed no synergism in expression levels between HFD and dexamethasone. This finding indicates that lipid accumulation in response to dexamethasone treatment is likely occurring via mechanisms other than accelerated glucocorticoid-dependent activation of *de novo* lipogenesis.

### Dexamethasone Causes Decreased Fat Mass in Obese Mice

To understand the how dexamethasone effects body composition in these animals, we determined total fat mass. We observed reductions in fat mass in the HFD-fed dexamethasone-treated group (Figure 3A-B). These reductions do not appear to be depot-specific, as we observed reductions in both iWAT (65% reduced) and eWAT mass (59% reduced; Figure 3C) ub the obese, dexamethasone-treated mice. There were no significant differences in fat mass, either by MRI or gross tissue weights of iWAT or eWAT depots in response to dexamethasone treatment in the chow-fed groups (Figure 3B-C). To determine if changes in body composition could be explained by altered caloric consumption (Figure 3D), we compared food intake among the groups. Chow-fed, dexamethasone-treated mice ate significantly less than chow-fed controls (9% reduction; p=0.006), as previously reported (Haber & Weinstein 1992; Roussel *et al.* 2003). Surprisingly, we found that the dexamethasone-treated HFD-fed animals ate slightly more food (11% increase, p=0.032), even though they lost both fat and fat-free mass. These data suggest that the weight loss in obese animals provided dexamethasone is not due to reductions in food intake.

**Figure 3:**
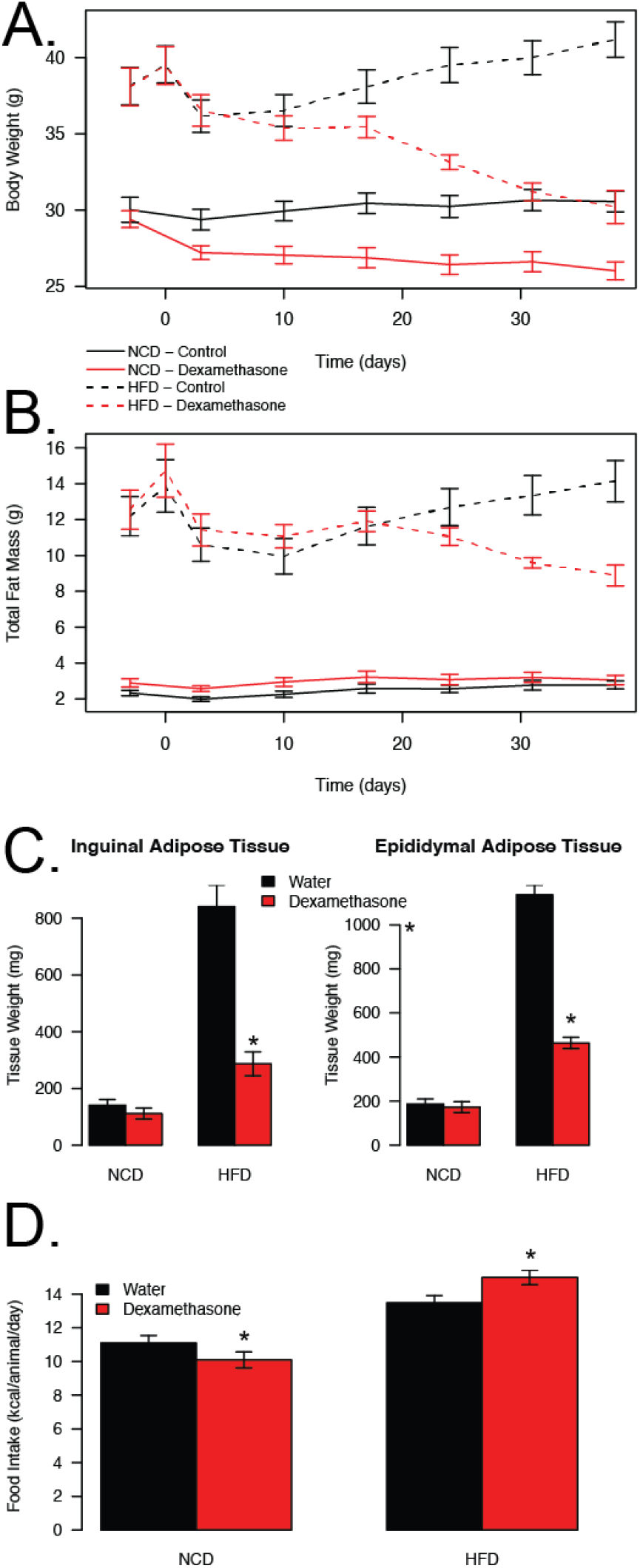
Dexamethasone treatment reduces fat mass in obese mice. Weekly total body mass (A) and fat mass (B) measures via EchoMRI in mice over the course of treatment (solid lines represent NCD mice and dashed lines represent HFD mice). Adipose tissue weights in 16 hour fasted mice following euthanasia (C). Mice were euthanized at 28 weeks of age following six weeks of dexamethasone (NCD n=8; HFD n=12) or vehicle (NCD n=8; HFD n=22) treatment and 18 weeks of diet. Food consumption measured weekly over the course of treatment (D). Asterisks indicate a statistically significant treatment effect for the pairwise comparison.

### Dexamethasone Treatment Results in Increased Lipolysis

Lipolysis has previously been associated with insulin resistance (Rebrin *et al.* 1996; Edgerton *et al.* 2017), is known to be elevated in patients with NAFLD (Gastaldelli *et al.* 2009), and has been shown to increase with high levels of glucocorticoids (Djurhuus *et al.* 2002, 2004; Kršek *et al.* 2006; Hochberg *et al.* 2015). To assess whether dexamethasone was affecting the lipid content in adipose tissue, we measured markers of adipocyte lipolysis in cultured adipocytes. 3T3-L1 fibroblasts were undifferentiated (pre-adipocytes); or differentiated and treated with vehicle or dexamethasone following differentiation. Dexamethasone treatment following differentiation led to decreased lipid content (52.4% reduction, p=0.005) and a 71% increase in the amount of glycerol in the media (p=0.001), suggesting increased lipolysis (Figure 4B). In order to identify a potential GR-dependent lipolytic target, we evaluated the levels of ATGL, the rate limiting enzyme in lipolysis. Expression of ATGL (encoded by the *Pnpla2* gene) was enhanced following dexamethasone treatment in 3T3-L1 cells at the transcript (2.7 fold, p=0.002; Figure 4C) and protein (4.2 fold, p=0.025; Figure 4D-E) levels. These data show that glucocorticoids elevate ATGL levels and metabolites of lipolysis in cultured adipocytes.

**Figure 4:**
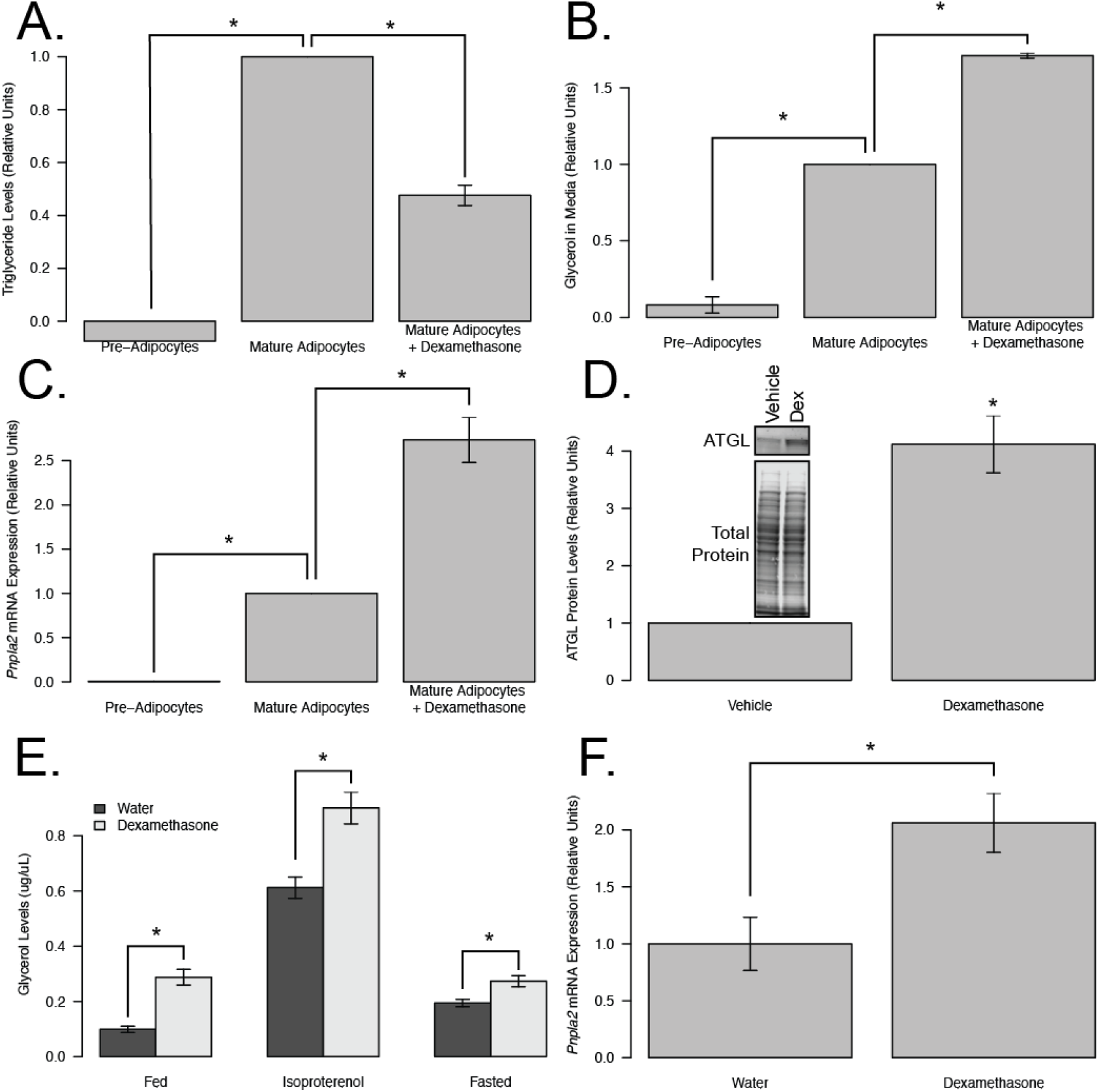
Dexamethasone treatment induces lipolysis in *vivo* and *in vitro*. Triglyceride levels (A), glycerol released in media (B), qPCR of lipolytic transcripts (C), and western blot of ATGL (D) from non-differentiated (pre-adipocytes; n=2) or differentiated 3T3-L1 mouse adipocytes (mature adipocytes) following five days of dexamethasone (n=3) or vehicle treatment (n=3). Serum fatty acid and glycerol levels at basal (fed) and following stimulation (10mg/kg isoproterenol or 16hr fast; E) and qPCR of IWAT lipolytic transcripts (F) in 22-week-old, 12-week dexamethasone-(basal and isoproterenol n=7; fasted serum and qPCR n=4) or vehicle-(basal and isoproterenol n=12; fasted serum and qPCR n=11) treated, chow-fed mice with the exception of isoproterenol-stimulated glycerol, which was performed one week prior to euthanasia. Asterisks indicated statistically significant treatment effect for the pairwise comparison.

To measure the effects of glucocorticoid-induced lipolysis *in vivo,* we quantified glycerol levels in animals chronically exposed to dexamethasone in basal and stimulated conditions (Figure 4E). Stimulation of lipolysis was achieved via isoproterenol, a β-adrenergic receptor agonist, or by a 16-hour fast. Dexamethasone treatment led to increases in glycerol in the fed (2.9 fold), fasted (1.5 fold) and isoproterenol-stimulated (1.4 fold) conditions (p<0.05 for all pairwise comparisons), indicating that dexamethasone enhances basal and stimulated lipolysis *in vivo* in chow-fed mice. Consistent with these findings, mRNA analysis from iWAT of these mice showed an upregulation of *Pnpla2* transcripts in the dexamethasone-treated mice compared to controls (2.1 fold, p=0.016; Figure 4F).

To understand how diet-induced obesity alters dexamethasone-induced lipolysis, we next quantified serum glycerol concentrations in our HFD/NCD fed mice (Figure 5A). We observed a nearly two-fold increase in serum glycerol levels by dex-amethasone treatment in the HFD-fed animals, compared with only a 18% increase in chow-fed mice (p=0.017 for the interaction between diet and dexamethasone). For the hyperinsulinemic euglycemic clamp in the obese mice there was a 40% elevation in serum basal non-esterified fatty acids (NEFA’s) in response to 3 weeks of dexamethasone treatment (p=0.004; Figure 5B). During the insulin phase, dexamethasone treatment attenuated the ability of insulin to suppress serum NEFA levels with insulin leading to a 71% reduction in controls compared to only a 48% reduction in dexamethasone-treated mice (p=0.058). These findings suggest that in the obese setting, dexamethasone elevates lipolysis to a greater extent and attenuates the effects of insulin.

**Figure 5:**
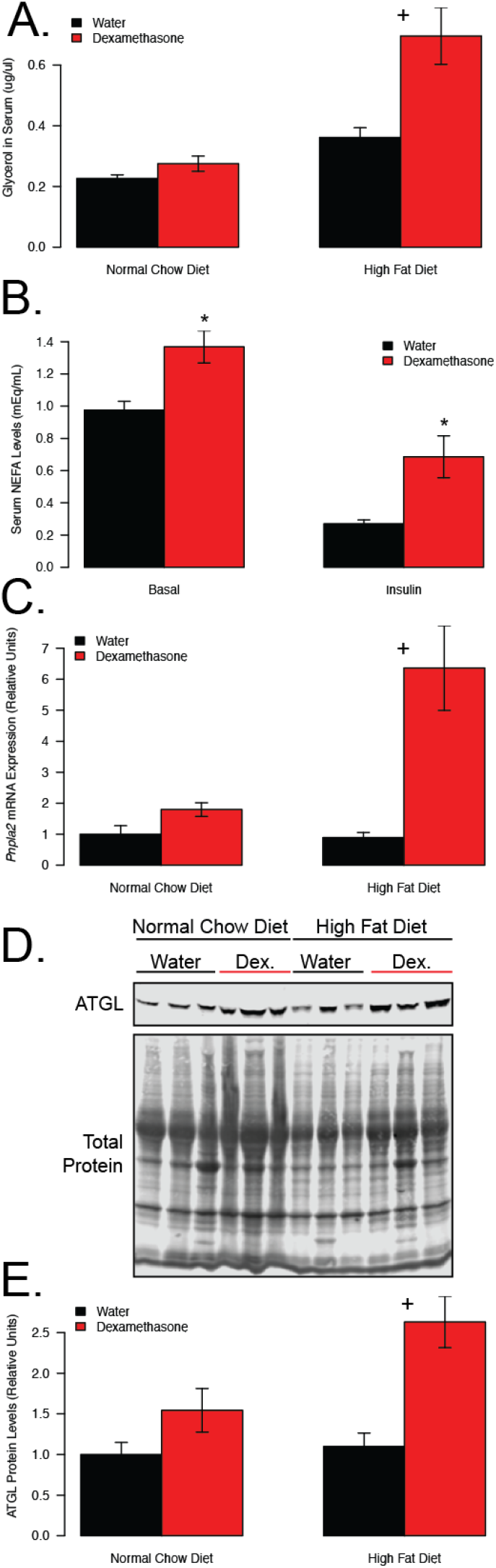
Obesity exacerbates dexamethasone-induced lipolysis. Serum glycerol (A) following 16 hour fast, serum NEFA in obese dexamethasone treated (n=14) or control (n=11) mice following a 5 hour fast, before and after insulin during hyperinsulinemic euglycemic clamp (B), qPCR of *Pnpla2* transcripts from iWAT (C), and western blot image (D) and quantification (E) of ATGL protein from iWAT. Mice from A, C, D and E were euthanized at 28 weeks of age following six weeks of dexamethasone (NCD n=8; HFD n=10) or vehicle (NCD n=8; HFD n=10) treatment. Crosses indicate a significant interaction between diet and treatment. Asterisks indicate a statistically significant treatment effect for the pairwise comparison.

To investigate the molecular basis for this synergistic increase in lipolysis, we quantified mRNA and protein expression of ATGL in the iWAT of these mice (5C-E). Consistent with the hypothesis that ATGL activation could drive increased lipolysis in obese dexamethasone-treated mice, expression of ATGL was elevated in both dexamethasone-treated groups, with a significant synergistic effect of dexamethasone and obesity at the transcript (p=0.02) and protein (p=0.043) levels. These data support the hypothesis that glucocorticoid-stimulated lipolysis is augmented in the context of obesity, potentially via increased transactivation of *Pnpla2*/ATGL.

## Discussion

Chronic glucocorticoid elevations are associated with co-morbidities such as increased fat mass (Abad *et al.* 2001; Geer *et al.* 2011; Hochberg *et al.* 2015), decreased muscle mass (Dardevet *et al.* 1995; Schakman *et al.* 2013; Hochberg *et al.* 2015), insulin resistance and NAFLD (Paredes & Ribeiro 2014). Many of these adverse effects are similar to those seen in obesity; however, the combination of chronically elevated glucocorticoids in the context of pre-existing obesity has not been assessed. Here, we show that the effects of glucocorticoid-induced insulin resistance and NAFLD are exacerbated when paired with obesity.

We appreciate that glucocorticoids directly affect many other tissues, such as muscle, liver and the pancreas that may also influence insulin sensitivity. In support of a central role of adipocytes, several studies demonstrate that adipocyte-specific reductions in glucocorticoid signaling being associated with improved metabolic profiles (Morgan *et al.* 2014; Wang *et al.* 2014; Mueller *et al.* 2017; Shen *et al.* 2017). We hypothesize that adipose tissue lipolysis plays a major role in dexamethasone-induced insulin resistance and hepatic steatosis, especially in the case of obesity.

Excess adiposity, such is seen in obesity, has been associated with increased insulin resistance. Previous work from our lab shows increased fat mass following 12 weeks of dexamethasone treatment (Hochberg *et al.* 2015) in lean mice, in accordance with what others have reported (Burke *et al.* 2017), as well as reduced insulin sensitivity. However, to our surprise, the glucocorticoid treatment in obese mice led to a lipodystrophic phenotype, which indicates the disturbances in glucose homeostasis are not a result of increased fat mass. The loss in fat mass observed in the obese, dexamethasone treated mice was not due to reduced food intake, in fact these mice ate significantly more calories per day than obese controls. This suggests a potential increase in energy expenditure with the combination of obesity and dex-amethasone treatment over time. This study evaluated glucocorticoid treatment in the context of diet-induced obesity; however, Riddell and colleagues have reported similar findings when providing HFD and glucocorticoids in concert to rats, prior to the onset of obesity (D’souza *et al.* 2012; Shpilberg *et al.* 2012; Beaudry *et al.* 2013). It is not clear whether diet or obesity status have similar mechanisms causing exacerbated metabolic risk, but these interactions deserve further evaluation.

Lipolysis has been linked to increased gluconeogenesis by several studies (Williamson *et al.* 1966; Nurjhan *et al.* 1986, 1992, Perry *et al.* 2015, 2017). Glucocorticoids are known to stimulate lipolysis (Djurhuus *et al.* 2002, 2004; Kršek *et al.* 2006; Hochberg *et al.* 2015), possibly as a way to promote gluconeogenesis to maintain blood glucose levels. Lipolysis has been implicated in insulin resistance (Rebrin *et al.* 1996; Edgerton *et al.* 2017) and NAFLD (Gastaldelli *et al.* 2009). We found synergistic elevations in glycerol, indicative of enhanced lipolysis, as well as in hepatic fat accumulation in the HFD-fed, dexamethasone-treated mice, but no data supporting enhanced hepatic *de novo* lipogenesis. Therefore, we propose the dexamethasone-induced increase in hepatic steatosis in the obese mice is primarily due to enhanced lipolysis observed in these animals.

There is some debate as to which genes glucocorticoids are acting on to promote lipolysis. Downregulation of *Pde3b* (Xu *et al.* 2009) and upregulation of β-adrenergic receptors (Lacasa *et al.* 1988) and ATGL transcripts (Campbell *et al.* 2011; Serr *et al.* 2011; Shen *et al.* 2017) have been proposed as possible mechanisms. We found ATGL, the rate limiting enzyme for adipose triglyceride lipolysis, to be synergistically activated by obesity and glucocorticoid-treatment. These findings bear a resemblance to elevations in glycerol levels in obese, dexamethasone-treated mice when compared to diet or glucocorticoids alone. The mechanisms by which obesity and glucocorticoids synergize to activate ATGL expression are not clear at this time, nor are the relative contributions of other glucocorticoid receptor-dependent targets.

In summary, glucocorticoids are commonly prescribed drugs used to treat a multitude of health issues, but are known to induce a variety of adverse metabolic effects. Their actions in persons with obesity are not yet clear, even though there is a significant number of individuals with obesity routinely taking prescription glucocorticoids. This paper is the first to show that diet-induced obesity in mice exacerbates several co-morbidities associated with chronically elevated glucocorticoids. These effects may be considered by physicians when determining glucocorticoid treatment options for patients with obesity.

### Declaration of Interest

The authors declared no conflict of interest that could be perceived as prejudicing the impartiality of the research reported.

### Funding

This study was supported by funds from NIH Grant R01-DK107535 (DB) and a Pilot and Feasibility Grant from the Michigan Diabetes Research Center (P30-DK020572). This study also utilized the University of Michigan Metabolism, Bariatric Surgery and Behavior Core (U2C-DK110768), the Michigan Nutrition Obesity Research Center (P30-DK089503) and the University of Michigan Comprehensive Cancer Center Core (P30-CA062203). Erin Stephenson is partially supported by funding from Le Bonheur Children’s Hospital, the Children’s Foundation Research Institute and the Le Bonheur Associate Board.

### Author contributions

D.B. acquired funding. D.B., I.Ha. and I.Ho. were responsible for conceptualizing the study. D.B., I.Ha. and N.Q. designed the experiments. I.Ha. performed all cell experiments. I.Ha., E.S. and J.R. performed mouse experiments. D.B. and Q.T. performed statistical analyses. I.Ha. wrote the manuscript. I.Ha. and D.B. edited and reviewed the manuscript. All authors were involved in discussions. This manuscript has been approved by all authors.

## Acknowledgements

We would like to thank the study participant for their willingness to be involved in this research. We would like to thank Jennifer Del-Proposto and Carey Lumeng for assistance with imaging liver sections, and Melanie Schmitt for assistance with glucose clamp studies. We would like to thank the other members of the Bridges laboratory, Thurl Harris (University of Virginia), Christoph Buettner and Eliza Geer (Icahn School of Medicine at Mount Sinai) and Edwards Park (UTHSC) for insights on this work. g

